# From Specification to Implementation: Assume-Guarantee Contracts for Synthetic Biology

**DOI:** 10.1101/2022.04.08.487709

**Authors:** Ayush Pandey, Inigo Incer, Alberto Sangiovanni-Vincentelli, Richard M. Murray

**Affiliations:** Control and Dynamical Systems at the California Institute of Technology; Electrical Engineering and Computer Sciences at the University of California, Berkeley

## Abstract

We provide a new perspective on using formal methods to model specifications and synthesize implementations for the design of biological circuits. In synthetic biology, design objectives are rarely described formally. We present an assume-guarantee contract framework to describe biological circuit design objectives as formal specifications. In our approach, these formal specifications are implemented by circuits modeled by ordinary differential equations, yielding a design framework that can be used to design complex synthetic biological circuits at scale. We describe our approach using the design of a biological AND gate as a motivating, running example.

## I. Introduction

Mathematical modeling has played a key role in the foundations of synthetic biology. In two papers published in 2000, dynamical systems were used to successfully analyze the oscillatory response of the repressilator [1] and the bistability in the toggle switch [2]. Since then, mathematical modeling has been extensively used to study the design of engineered biological systems [3]. Phenomenological models of gene regulation in synthetic biological circuits, such as activation and repression, have seen the most success in various applications [4]–[7]. These models use simple nonlinear functions to model the biological circuit phenomena representing their input-output behavior. Detailed models of mechanisms and metabolic pathways have long been used in systems biology to quantitatively study various properties of biological systems [8]. This kind of detailed modeling is important in engineered biological systems, as well. For example, chemical reaction network models of protease-based biosensors have been developed to engineer optimized logic gates [9]. Similarly, detailed mathematical models have been used to amplify biofuel production [10] and to characterize fluorescent protein maturation [11]. A review article on the current modeling practices in synthetic biology [12] provides further references on this topic.

The choice of using a detailed or a phenomenological model is made according to the problem at hand. From a design standpoint, the development of models is necessary to decide when to use the circuits described by these models. As system complexity increases, we believe that it is necessary to develop a complete design methodology that begins with a top-level description of the system’s objective and guides the designer in the generation of an implementation that can be proved to meet the specification. This is the main contribution of this paper. Our methodology decouples reasoning about component specifications from reasoning about the modeling details of each component. This allows understanding the system’s specification by analyzing the specifications of the subsystems. This methodology allows designers to focus on particular aspects of the design process at various levels of detail while ensuring that other aspects of the design are not forgotten. Current design approaches in synthetic biology do not model system specifications mathematically, but rather informally. As a consequence, the resulting mathematical models obtained are often disconnected from the specifications. It is thus difficult to answer whether the obtained models behave according to the specifications.

The design methodology presented here is centered on contract-based design [13]. At the heart of contract-based design are assume-guarantee contracts, which are formal specifications that distinguish between the responsibilities of a component and the assumptions made on its environment. Contracts come with a rich algebra. Contract-based methodologies have been developed for digital circuit design, aircraft power distribution systems [14], and certain classes of control systems [15]. We demonstrate the application of our methodology with the design of a biological AND logic gate. We use contract algebra to derive the specification of the system from the specifications of its components. We discuss how contracts help us to meet a system-wide specification when we have a partial implementation of the system available. Finally, we show how our methodology seamlessly connects the specification of a component with its models and implementations.

### Organization

Section II describes the contract-based design methodology and the algebraic aspects of contract specifications. Section III applies the methodology to the design of a biological AND gate. A discussion follows. The appendix provides details of logical manipulations of contracts.

## II. A contract-based design primer

Contract-based design [13] is a methodology by which a system is implemented by starting from a specification expressed as a contract and proceeding through a process of successive refinements, each of them adding increasing amounts of detail to the implementation.

The design process begins with a contract for the system we wish to implement and with a library of components that will be used to construct such a system. Each component in the library is represented by a contract. A mapping process identifies a set of elements from the library whose composition implements the top-level contract. If an implementation cannot be found, it may be the case that the library is insufficiently populated to implement the given specification. In this case, we can identify the contract specification of an element we need to add to the library to meet the specification. On the other hand, when an implementation is obtained by the mapping process, we have either reached our goal, or we may need to determine additional details of the candidate implementation. In the latter situation, the result of the mapping process can become the “top-level” contract input of the next mapping step that adds more details to the implementation. In other words, the design process can repeat as many times as needed until all details of the design are determined.

In contract-based design, the library of components consists of contracts; hence, we have to characterize as a contract every component we wish to add to our library. Since there are various kinds of analyses we may wish to perform on the system, there can be multiple contracts that can be associated with a given design element. For example, we may want to keep separate the functionality specification from the performance specification. The multiple angles from which we can look at components are called *viewpoints*.

Operating on formal specifications offers the advantage that we keep separate the purpose of the system from its implementation details. This allows the designer to carry out analysis at the specification level, without getting distracted by detailed models. Moreover, by keeping separate the various viewpoints of the design, the designer can focus on a given type of analysis without worrying about the specifics of other aspects.

### A. Formal aspects of assume-guarantee contracts

Assume-guarantee contracts are formal specifications that distinguish between (i) assumptions made on the environment and (ii) responsibilities attributed to the object being specified when it operates in an environment that meets the assumptions of the contract. An assume-guarantee contract 𝒞 is thus a pair (*a, g*) of constraints denoting assumptions and guarantees, respectively. We say that a component is an environment for the contract if it meets the constraints *a*; we say a component is an implementation for a contract if it meets the guarantees *g* provided it operates in an environment for the contract, i.e., if it meets the constraint *a* → *g*, where the arrow is logical implication (i.e., *a* → *b* = *a* ∨ ¬ *b*). For a treatment of assume-guarantee contracts and their algebraic aspects, see [16]–[18].

We assume that constraints come from a Boolean algebra, i.e., there are well-defined notions of conjunction, disjunction, and negation for constraints. The Boolean algebra of constraints generates on contracts a partial order called refinement. We say that 𝒞= (*a, g*) refines 𝒞′ = (*a*′, *g*′) (or that 𝒞′ abstracts 𝒞) when the environments of 𝒞′ are environments of and the implementations of 𝒞 are implementations of 𝒞′, i.e., when *a*′ ≤ *a* and *a* → *g* ≤ *a*′ → *g*′. We say that two contracts are equivalent if they have the same environments and the same implementations. Note that any contract (*a, g*) is equivalent to the contract (*a, a* → *g*). A contract in this form is said to be in saturated or canonical form.

Suppose we have two components obeying contracts 𝒞 and 𝒞′, respectively, and we want to obtain the specification of the system built using these two components. The contract operation of *composition* gives us the smallest contract in the refinement order obeyed by the system. Its closed-form expression is

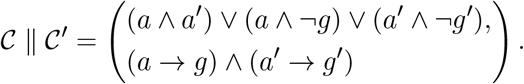

The composition operation is monotonic in the refinement order, i.e., composing by bigger contracts yields bigger results. As the composition is the smallest contract obeyed by the system, the system also obeys any abstraction of the composition operation. This observation will be used to provide results of composition that are closer to the intuition of a designer.

In addition to building systems using components, we are sometimes interested in finding components that allow us to meet a goal system-level specification. Suppose we wish to implement a system having a specification 𝒞, and we have available a subsystem obeying a contract 𝒞′.

We want to find the specification of a second subsystem whose composition with the existing subsystem satisfies the top-level specification 𝒞. The largest such contract in the refinement order is given by the operation of *quotient*, whose explicit form is

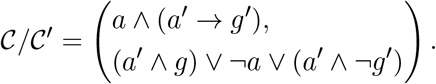

Since the quotient is the biggest specification that completes the system, any contract smaller than the quotient also completes the system. As before, we will find it convenient to refine the quotient to provide contracts that match the intuition of the designer.

In addition to composing and decomposing systems, contracts offer support for multi-viewpoint design [19]. As we discussed, we are also interested in carrying out analysis of systems by focusing on one aspect of the system. This aspect may be functionality or performance. This means that we can assign to each component in our system several contracts, one for each viewpoint. The operation of *contract conjunction*, or *weak merging*, can be used to summarize into a single contract two viewpoints of the same object. Conjunction is given by the expression

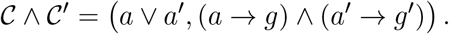

## III. Contract-based Design in Synthetic Biology

In this section, we consider the use of contracts to compose and decompose biological systems. First, we provide an overview of the system we are constructing, a biological AND gate. We introduce the contracts of the components comprising the system using two viewpoints: one for functionality and one for timing. Then we compose the contracts to obtain the specification of the entire system by using the properties of the components. We compose the models of each contract to verify that this composition meets the top-level specification we obtained. Then, starting from a top-level specification of the system and the specification of a partial implementation, we obtain the specification of a missing component that enables us to meet the top-level specification. We synthesize a model from this specification, and we show that the resulting composition of models implements the top-level specification.

### A. Biological AND gate

Consider the design of an AND gate with inputs *u*_1_ and *u*_2_ and one output *y*. A biological circuit design to achieve the AND logic has been built and validated in [20]. This design is a composition of three subsystems that interact, as shown in Figure 1. In [20], the inputs *u*_1_ and *u*_2_ are chemical inducer signals (salicylate and arabinose respectively), and the output *y* is the fluorescence of green fluorescent protein (GFP). Let the output of subsystem Σ_1_ be *x*_1_ that models the expression of an amber suppressor tRNA called supD in [20]. Similarly, the output *x*_2_ of Σ_2_ models the transcription of the mRNA that codes for the T7 RNA polymerase enzyme. However, the translation of the mRNA (*x*_2_) is only possible when *x*_1_ is also present. When both *x*_1_ and *x*_2_ are present, the translation of the T7 RNA polymerase can occur, which activates the T7 promoter in Σ_3_ that codes for a fluorescent protein GFP, the output *y* of the system. The top-level specification for the AND gate is given in Table I.

**TABLE I:** The static AND gate specifications are such that when both inputs *u*_1_ and *u*_2_ are greater than their specified minimum values, we have *y*^*^ ≥ *Fy*_*ϵ*_, where *F* > 1 is the desired fold change in output compared to the leaky output *y*_*ϵ*_. The dynamic specifications add that the output achieves the desired fold-change in time *t* ≥ *τ*_*y*_.

| $u_1$ | $u_2$ | $y$ | For $u_1 \geq u_{1\min}, u_2 \geq u_{2\min}$: |
| --- | --- | --- | --- |
| 0 | 0 | $y_\epsilon$ | |
| $u_{1\min}$ | 0 | $y_\epsilon$ | |
| 0 | $u_{2\min}$ | $y_\epsilon$ | |
| $u_{1\min}$ | $u_{2\min}$ | $y^*$ | $y^* \geq Fy_\epsilon, \quad t \leq \tau_y$ |

**Fig. 1:**
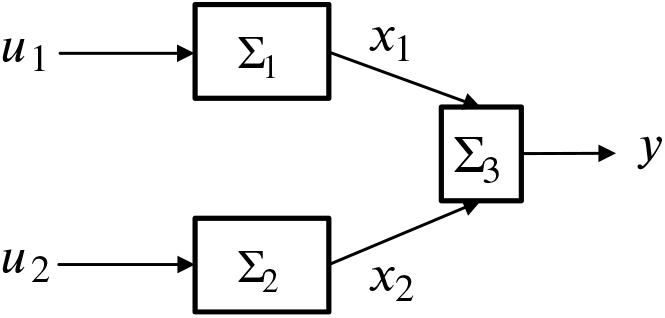
Composition of three subsystem models to achieve AND logic gate implementation in an engineered biological system.

We can write the specification for the subsystem Σ_1_ by describing the assumptions and guarantees of the biological design. We assume that at time 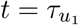 we have 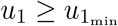, and *u*_1_ stays over this threshold. The contract 𝒞_1_ = (*a*_1_, *g*_1_) for Σ_1_ guarantees that 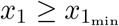 at time 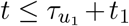. Here *u*_1_ is the salicylate inducer concentration, and *x*_1_ is the expression level of the supD gene that is downstream of the pSal promoter.

We split our specification in two viewpoints. First, we have a functionality viewpoint that says that 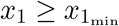 follows from 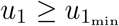. The other is a timing viewpoint that says that the event 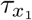, defined as the time when 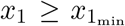, happens at most *t*_1_ time units after the event 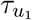, defined as the time when 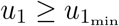. That is, we have the following two contract viewpoints:

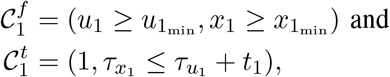

where 1 in the assumptions of the last contract represents the top element of the constraint lattice, i.e., the boolean value “true.”

For Σ_2_, we have the input *u*_2_ (arabinose) that activates the pBAD promoter to express the T7Ptag gene downstream. For this subsystem, if we assume that at 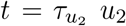 crosses the threshold 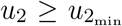, then the subsystem specification guarantees that 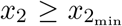 at time 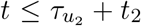. The functionality and timing contracts *C*_2_ = (*a*_2_, *g*_2_) for Σ_2_ are

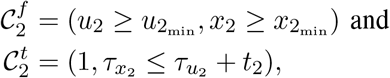

where 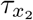 is, as before, the event when *x*_2_ crosses the threshold 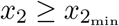.

For Σ_3_, we have inputs *x*_1_, the tRNA supD, and *x*_2_, the engineered T7 RNA polymerase transcript from Σ_1_ and Σ_2_ respectively. The translation of T7 RNA polymerase occurs only in presence of both *x*_1_ and *x*_2_ that then drives the production of the output, *y*, the green fluorescent protein. Under the assumptions that 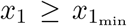 and 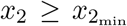 starting at some 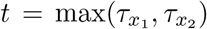, Σ_3_ guarantees that the output *y* is at least *F* > 1 fold-change higher than the leaky expression output *y*_*ϵ*_ at time 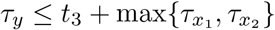. Hence, the contracts for Σ_3_ are

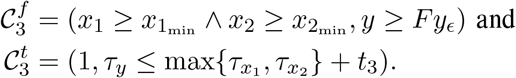

### B. Generating specifications of the system

Now that we have the specifications for the three elements of the system, we seek the specification of the entire system. First, we use the operation of composition to obtain the specification of the subsystem consisting of components Σ_1_ and Σ_2_. We compute 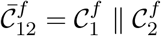 and 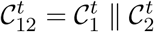:

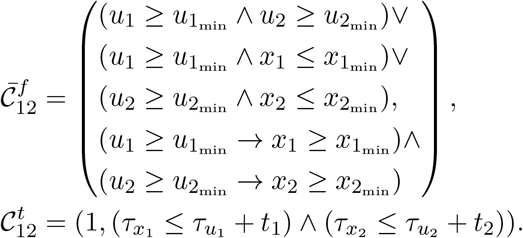

Composition yields the smallest specification that the system obeys. This means that the system satisfies any specification which is more abstract than that obtained from the composition operation. We observe that the second and third terms in the assumptions contain “failure terms,” i.e., terms in which the assumptions of components were met, but where the guarantees failed to be met. We eliminate these terms to obtain the abstraction

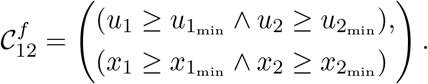

Now we obtain the specification of the entire system by composing these contracts with those of Σ_3_, i.e., we compute 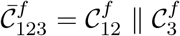 and 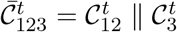 and then abstract these contracts to obtain 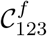and 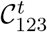:

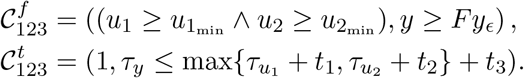

These contracts give us a specification for the entire system. They only refer to variables that lie at the interface between the system and its environment, namely *u*_1_, *u*_2_, and *y*; there is no mention of *x*_1_ and *x*_2_. This allows us to “black-box” the system so that it can be used as a component of a more involved system. Appendix A shows in detail the logical manipulations that allow us to compute the abstraction 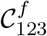.

### C. From specifications to dynamical models

We explore the link between the contract specifications and the component models as differential equations. As discussed above, the formal specifications describe the desired functional and timing objectives for the system. By developing dynamical models with biological details from these specifications, we can map the desired objectives to system implementations. We first develop first-order dynamical models by employing standard functions to convert time delays and function gain to dynamical equations. Then we add more detail to these models. A flow chart summarizing this process is shown in Figure 2. For Σ_1_, we can write the following equation:

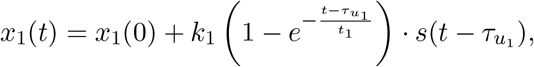

where *s*(·) is the step function such that for 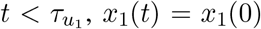and for 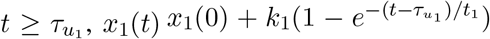. The parameter *k*_1_ is given according to the specification in the contract as

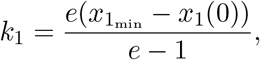

so that at 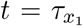, we have 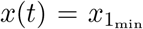. Note that this is an implementation of the contract 𝒞_1_ since it guarantees that 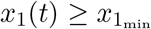 when the assumption 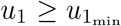 is satisfied. Further, the timing contract is also satisfied as the implementation guarantees that 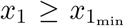 at time 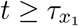. The input is modeled as an ideal activator function by using the step function. Following a similar approach for Σ_2_, we have

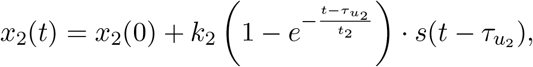

where

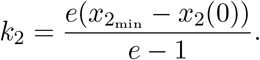

**Fig. 2:**
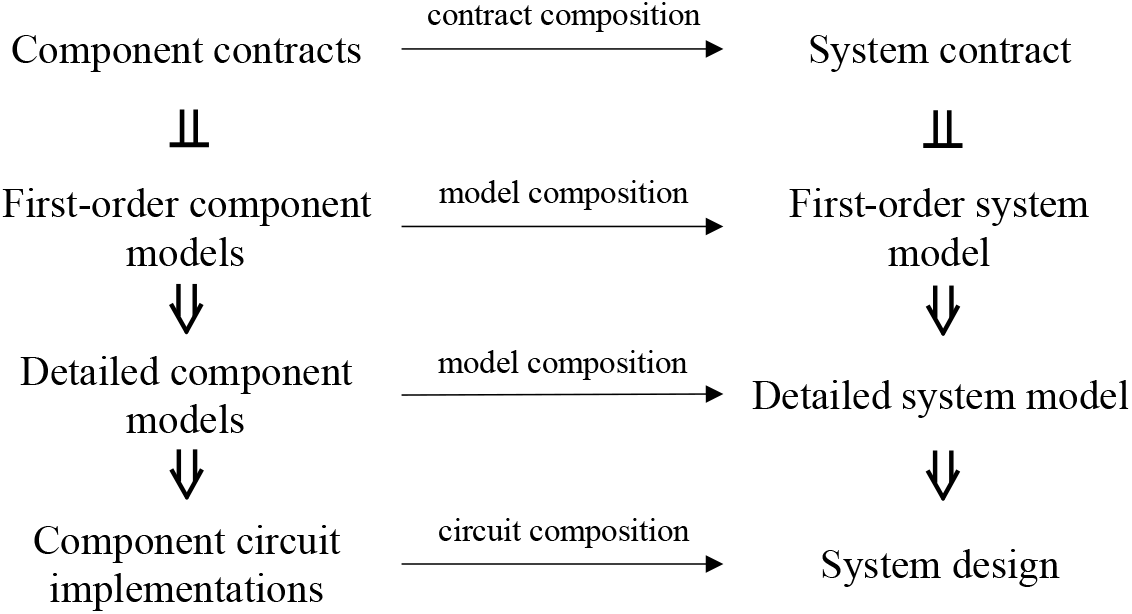
Contract-based design allows us to compute a system specification from the specifications of the system’s components. The system-level contract has the property that its implementations are the compositions of the implementations of the component contracts. This property holds even if we keep adding detail to our models.

Finally, for Σ_3_, define 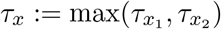. We have

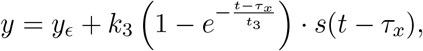

where

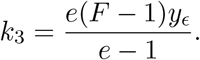

We now derive differential equations that will enable us to make a clearer connection between the first-order models and models that are more descriptive of biological implementations. We differentiate the equations for *x*_1_, *x*_2_, and *y* with respect to time to obtain

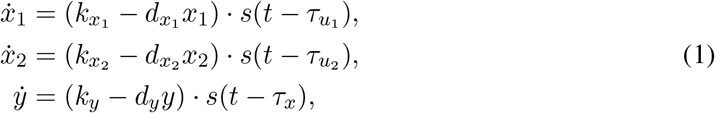

where

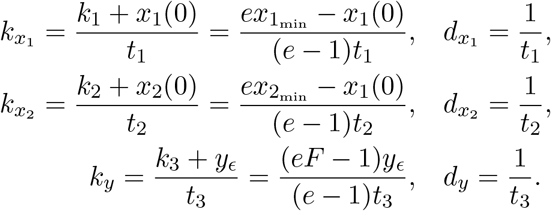

In the ODE model, for 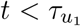 we have 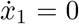, so we set the boundary condition as *x*_1_(*t*) = *x*_1_(0) for all 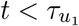. Similarly, we write the initial conditions as boundary values for *x*_2_(*t*) and *y*(*t*).

*Remark:* The dynamical model shown above follows from the component specifications written as contracts. Hence, the parameters in the models are functions of the component specifications. Numerical simulations of the dynamical models are shown in Figure 3. The simulations show that the composition of all system models is a valid implementation of the system-level contract specification we computed.

**Fig. 3:**
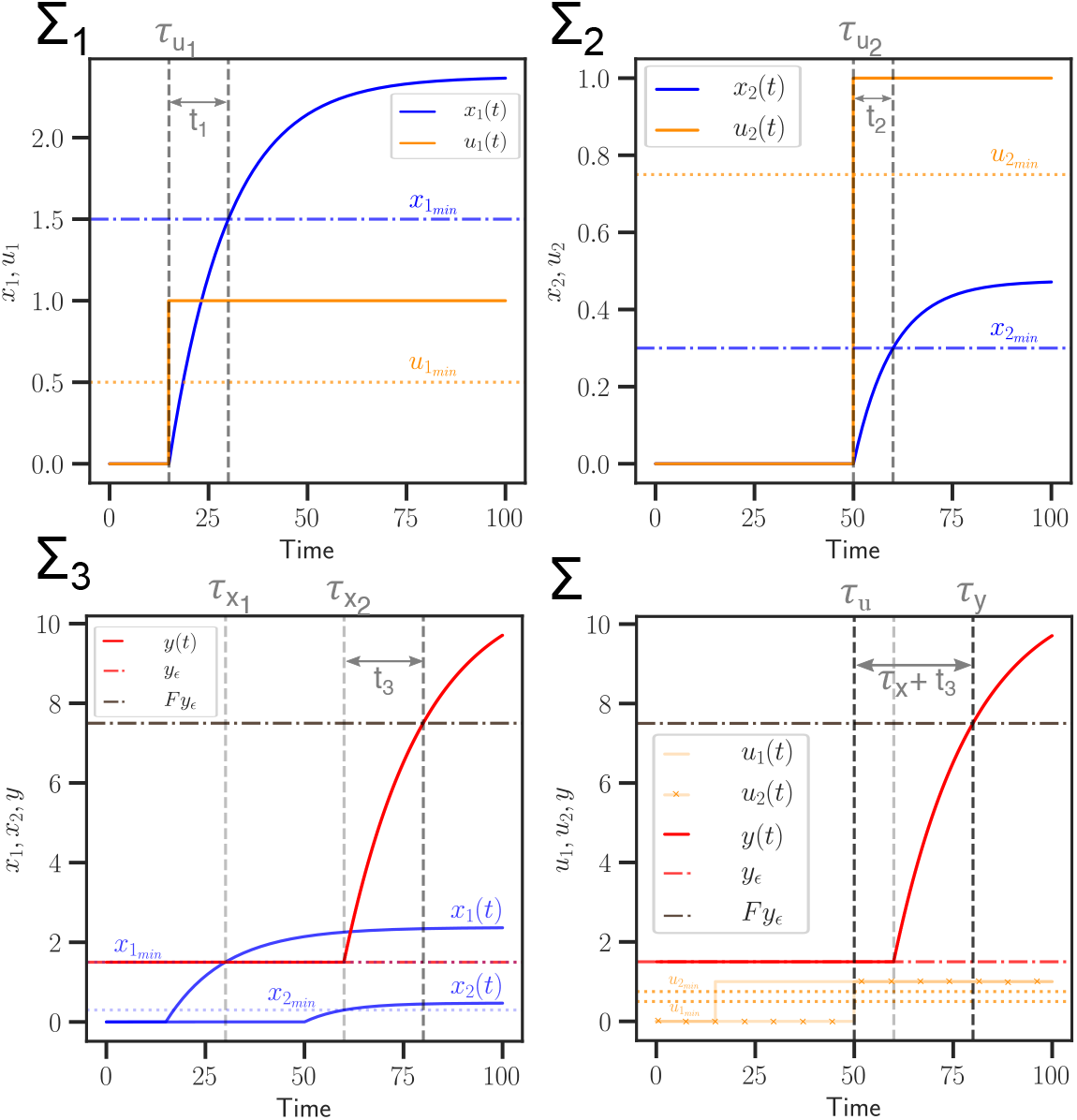
Simulations for each subsystem, Σ_*i*_, *i* = 1, 2, 3 and the composed system Σ are shown in the figure. Dotted and dashed lines represent the assumptions and guarantees, respectively, while the solid lines show an implementation of the contract. Python code to generate the plots is available online [21].

Observe that in the first equation of (1), for all 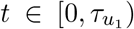, we have 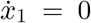 and for all 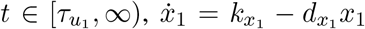. If the initial condition *x*_1_(0) = 0, then we can simplify the differential equation to 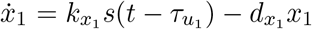 since the step function multiplied with the linear degradation term is redundant. We can follow a similar procedure for all equations in (1) to write ODE models such that the inputs to each subsystem activates its response through the production term. Under the assumption that the initial conditions are *x*_1_(0) = *x*_2_(0) = *y*(0) = 0, we have

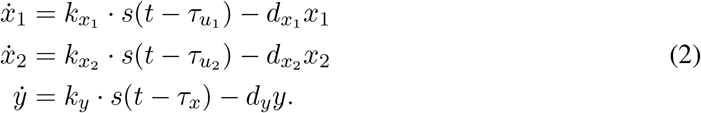

The composition of subsystems gives us an implementation of the top-level system contract as

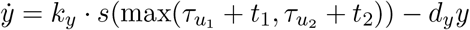

by using the contract definitions.

We have shown the synthesis of dynamical models for the subsystems and their composition from the specification contracts. However, these first-order models are not descriptive of biological implementations. As a result, it is not possible to map the design objectives to the controllable parameters in the implementation. In order to expand the model to include details of biological mechanisms, we observe that the first-order models use an ideal activation function — the step function. A common activation function used in the mathematical modeling of biological systems is a Hill function. An activation Hill function (rate = *u*^*n*^/(*u*^*n*^ + *K*^*n*^)) models activation as slow growth (rate ≪ 1) until a threshold (*K*) is reached, then, an ultra-sensitive response to asymptotically reach the maximum value (rate = 1). From our development of the mathematical models shown above, we can use a Hill function in place of the step function to model the non-ideal activation characteristics of an activatable promoter in the genetic circuit. An expanded model can be written as

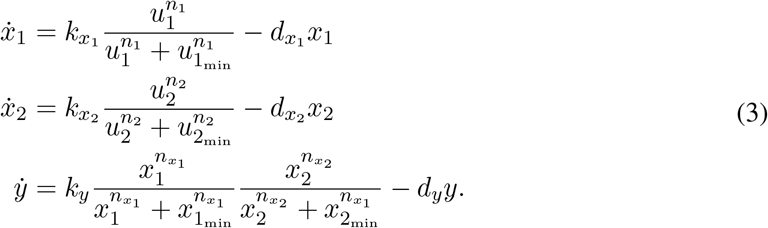

Note that in the limit of *n*_1_ → ∞, *n*_2_, and 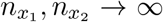, the respective Hill functions converge to a step function with the threshold *τ* variables defined according to the activation constants specified in the design (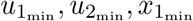, and 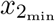). Hence, the model in (3) converges to the dynamical model in (2).

*Remark*: For a biological implementation, Hill function coefficients are usually constrained to *n*_*i*_ ≤ 4, where *i* denotes the subscripts for *n* above. With this additional constraint, the Hill function model may not meet the guarantees even when the assumptions are satisfied. We can offset this by letting the first-order model satisfy the guarantees such that *y* > *Fy*_*ϵ*_ at *t* ≪ *τ*_*x*_ +*t*_3_ so that even with a limited Hill coefficient, the detailed model can satisfy the stated guarantees in the contract.

Finally, we can expand the model in (3) to a chemical reaction network (CRN) model of the circuit implementation by using model reduction techniques such as conservation laws, state transformations, and time-scale separation, as discussed in [22]–[25] and related papers. The expansion to CRN models is out of scope for this work and maybe addressed in future research.

### D. Design synthesis of missing subsystem

In the previous two subsections, we went from component specifications to the specification of the entire system. Now we will start from a top-level specification and the specification of a subsystem, and we will look for the specification of the missing subsystem needed to satisfy the top-level specification. Therefore, suppose that we are given a specification for the entire system:

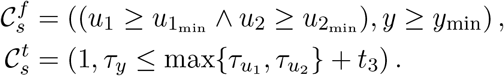

Suppose we also have available the specification of a subsystem, say the composition of Σ_1_ and 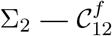 and 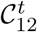 as computed above. The question is, what is the specification of an element that we have to add to 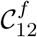 and to 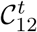 so that the resulting implementation meets the system-level specifications, 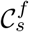 and 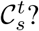 The largest specification with this property is given by the contract quotient. We compute the quotient 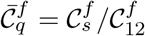 and 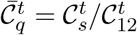:

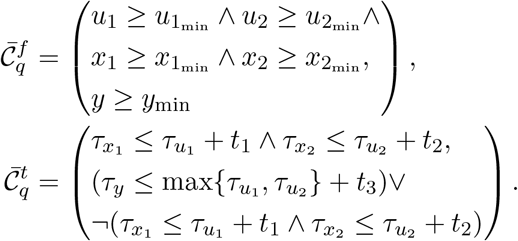

We can refine these two contracts by removing references to the inputs *u*_1_ and *u*_2_:

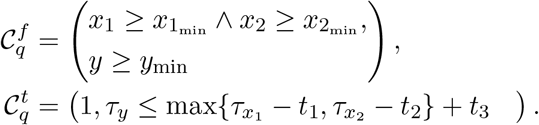

Observe that any implementation of this contract is guaranteed to satisfy the system-level specification when it operates in conjunction with an implementation of *𝒞*_2_. We now look for an implementation of *𝒞*_*q*_. We can propose a first-order model using the following expression:

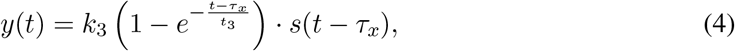

where we define 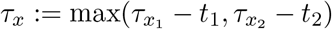 and

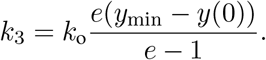

The dynamical model is given by

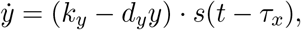

where

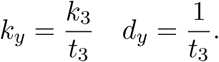

With *k*_o_ = 1, this model satisfies the guarantees such that *y* = *y*_min_ at the required timing guarantee *t* = *τ*_*x*_ + *t*_3_. This synthesized dynamical implementation model can be expanded further to include the modeling details specific to a synthetic biology implementation by using a Hill activator function (similar to model in (3)) instead of the step function. As discussed in the previous section, the biological implementation with a Hill function has an additional constraint on the Hill coefficient. To offset this, we set *k*_o_ > 1 so that the model in (4) satisfies *y* > *y*_min_ at a time *t* < *τ*_*x*_ + *t*_3_. In this way, the biological implementation with the Hill functions also satisfies the guarantees as shown in Figure 4.

**Fig. 4:**
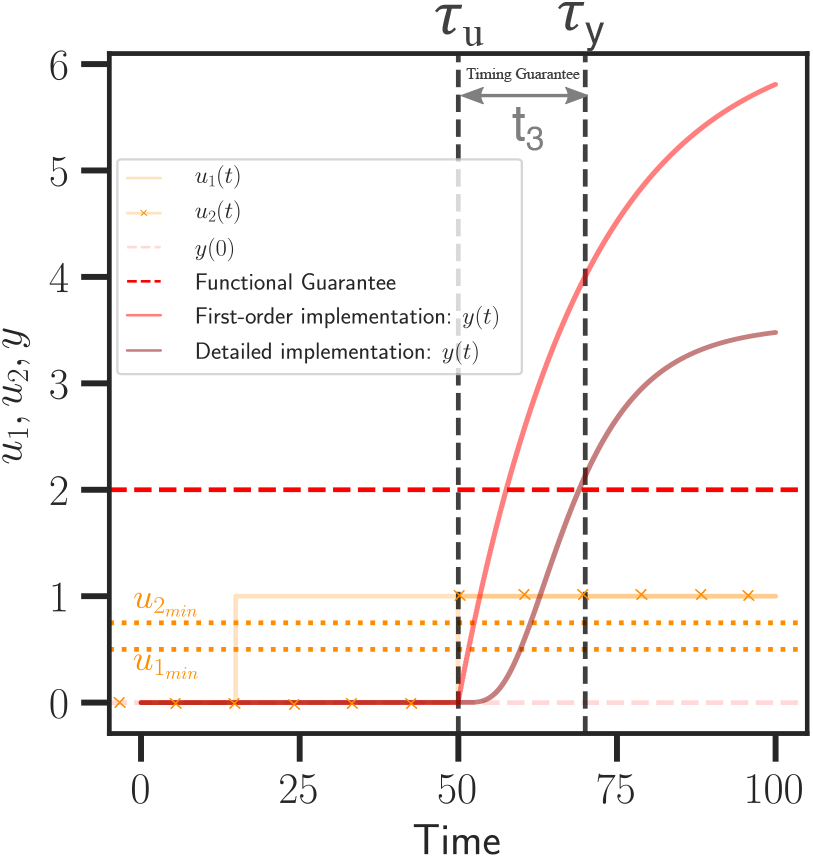
Synthesis of missing subsystem using the quotient operation of contracts. The dotted and dashed lines show the contract assumptions and guarantees, respectively, while solid lines show implementations. Here *τ*_*u*_ is defined as 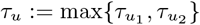.

## IV. Concluding remarks

### Summary

We presented a contract-based design framework for synthetic biology. Current modeling practices in synthetic biology are limited to system analysis and inverse problems for system identification. Our results are a step towards a design framework that reasons about system properties using contracts and is capable of correlating implementations with specifications. We showed how we could reason about system-level properties by composing the contracts of the components; then we composed the dynamical models for each component and verified that their composition satisfied the system-level contract. From the dynamical implementations, we derived detailed models that are closer to a biological implementation. Two key insights are discussed below.

### Speeding up the experimental design process

Usually, biological circuit design entails multiple experimental iterations to nail down controllable parameters such as promoter strengths, input levels, ribosome binding strengths, and physical conditions like pH and temperature. In our approach, since the parameters of the implementation models are mapped to the system objectives, these controllable aspects in the experimental design can be manipulated accordingly. Hence, a formal design framework serves to minimize the experimental trial-and-error steps required to successfully design a biological circuit. Moreover, we can analyze how uncontrollable parameters corresponding to the internal biological mechanisms in the implementation relate to the system objectives.

### Resource loading effects on system design

Engineered circuits with gene expression are dependent on cellular resources such as RNA polymerase, ribosome, ATP, and nucleic acids. For each subsystem to function as desired, a minimum level of these resources is required. At the same time, due to the resources being used by a subsystem, loading or retroactivity effects are commonly observed experimentally. Various system analysis approaches in the literature have analyzed retroactivity using mathematical models [26]–[28]. The contract-based design framework that we proposed in this paper can be easily scaled to describe such environmental assumptions as well. For example, consider RNA polymerase, ribosome, and ATP as three resources required for gene expression. We can add a resource viewpoint, 𝒞^*r*^, to the contracts for the subsystems in addition to the timing, 𝒞^*t*^, and the functionality, 𝒞^*f*^, viewpoints. The resource viewpoint contract for subsystem Σ_1_ can be written as

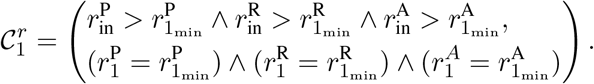

This contract states that if the input resources for polymerase 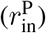, ribosome 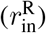, and ATP 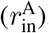, exceed minimum thresholds 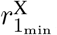, then the component Σ_1_ will consume a given amount of said resources 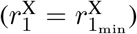. We can use this viewpoint to analytically detect when our system may not function due to starvation from a given resource. This viewpoint also models the amount of loading each subsystem causes to other subsystems. We anticipate diverse future directions stemming from the research presented in this paper as we build towards model-guided design for large-scale synthetic biological systems.

## Appendix

We consider the details of the derivation of 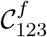. This contract is defined as an abstraction of 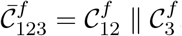, which we compute first by applying the definition of contract composition:

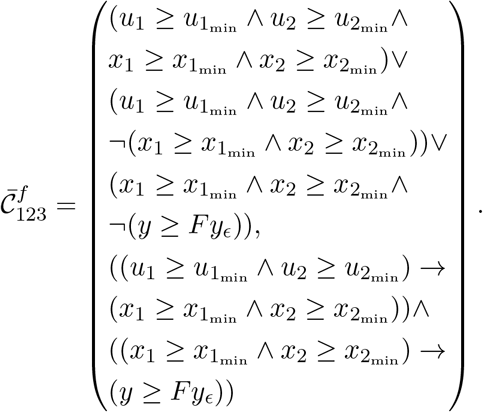

We focus on the first two terms of the assumptions. Observe that

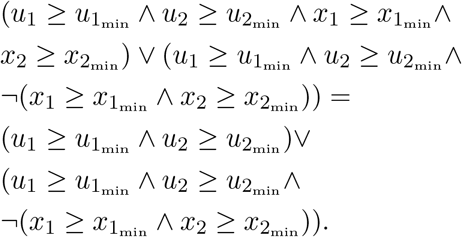

This simplification follows from the Boolean algebra identity *p* ∨ ¬*q* = (*p* ∧ *q*) *q*. After carrying out this simplification, we discard the second and third terms of the assumptions. Doing this yields smaller assumptions, which means that we obtain a contract abstraction. After carrying out this process, we obtain

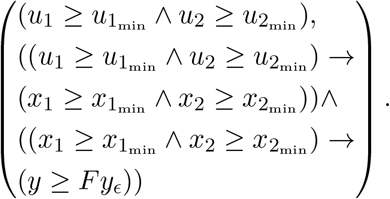

We place the guarantees of the contract in canonical form and carry out some simplifications:

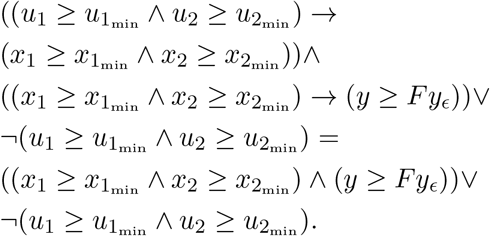

We can remove from the first term the terms involving *x*_1_ and *x*_2_ since these are internal variables that we not wish to include in the final system specification. Observe that removing terms from a conjunction yields bigger guarantees. We thus obtain a contract abstraction by carrying out this procedure. Finally, we hide the saturation term to produce the contract

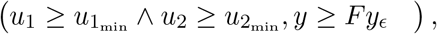

which is the contract 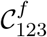.

